# Diet-based assortative mating through sexual imprinting

**DOI:** 10.1101/338848

**Authors:** E.K. Delaney, H.E. Hoekstra

## Abstract

Speciation is facilitated when traits subject to divergent selection also contribute to non-random mating—so-called ‘magic traits.’ Diet is a potential magic trait in animal populations because selection for divergence in consumed food may contribute to assortative mating and therefore sexual isolation. However, the mechanisms causing positive diet-based assortment are largely unknown. Here, using diet manipulations in a sexually imprinting species of mouse, *Peromyscus gossypinus* (the cotton mouse), we tested the hypothesis that sexual imprinting on a divergent diet could be a mechanism that generates rapid and significant sexual isolation. We provided breeding pairs with novel garlic- or orange-flavored water and assessed whether their offspring, exposed to these flavors *in utero* and in the nest before weaning, later preferred mates that consumed the same flavored water as their parents. While males showed no preference, females preferred males of their parental diet, which generated significant sexual isolation. Thus, our experiment demonstrates that sexual imprinting on dietary cues learned *in utero* and/or postnatally can facilitate reproductive isolation and potentially speciation.

## Introduction

The evolution of new species is easier when a trait undergoing divergent natural selection also causes assortative mating. The list of so-called “magic traits” – named for their seemingly magic effects on both adaptation and non-random mating (Gavrilets 2004) – is ever-growing, and includes examples of body size, color, and feeding morphology (Servedio et al. 2011). Diet has been recognized as a potential magic trait because of its likelihood of being under divergent natural selection between populations and because of its impacts on mate choice (Servedio et al. 2011). While diet-based assortative mating has been identified in multiple laboratory populations of fruit flies (Dodd 1989; Sharon et al. 2010) and natural populations of fish (Snowberg and Bolnick 2008, 2012; Martin 2013; Colborne et al. 2016), its cause is less well studied. Thus, asking how diet-based assortative mating arises is an important question in speciation research.

Assortative mating based on diet could arise if individuals select mates based directly on their diet, indirectly on traits correlated with diet, or incidentally based on non-heritable nutritional condition (Rosenthal 2017). Most studies examining mechanisms for diet-based mate choice have been limited to *Drosophila* and fish. In *Drosophila*, assortative mating preferences by diet were proposed to result from correlated dietary traits. It was suggested that feeding flies of the same strain different diets significantly altered their gut microbiota, changing pheromone mating signals as a result (Sharon et al. 2010; Rosenberg et al. 2018). However, such patterns of diet-based assortative mating were only found in inbred, not outbred, *Drosophila* strains (Najarro et al. 2015; Leftwich et al. 2017, 2018), calling into question the relevance of diet-based assortative mating and microbe-mediated pheromone effects in natural populations. In threespine stickleback and Cameroon crater lake cichlid fishes, diet-based assortative mating appears to be partially due to active mating preferences for diet or correlated traits (Snowberg and Bolnick 2012; Martin 2013); however, it is still unclear how individuals use dietary information to select mates.

We propose that sexual imprinting could provide a missing mechanistic link between diet and mate choice. That is, in species with parental care, offspring might learn to prefer the diet of their parents, leading to sexual isolation when mates are selected based on diet. Dietary information could be conveyed visually (e.g., diet-derived pigments, such as carotenoids) or through chemical odors and pheromones (e.g., potentially mediated through the gut microbiome). For example, changes in diet have been shown to alter individual body odors (Ley et al. 2008) and affect pheromone production or metabolites in rats, swordtails, and fruit flies (Leon 1975; Bell et al. 1991; Phipps et al. 1998; Fisher and Rosenthal 2006; Sharon et al. 2010). Should a source of divergent natural selection favor a shift in individual diets, sexual imprinting on detectable dietary cues during a sensitive period either *in utero* or shortly after birth could generate diet-based assortative mating.

Here, we experimentally test the hypothesis that changes in diet, when sexually imprinted, will lead to assortative mating. We first manipulated diet in cotton mice *(Peromyscus gossypinus*) – a species in which individuals are known to sexually imprint on their parents (Delaney and Hoekstra 2018) – by providing breeding pairs either garlic- or orange-flavored water. We then tested if offspring preferred mates fed on the same diet as their parents, thereby creating diet-based assortative mating. We present results that show sexual imprinting on diet is possible and can lead to assortative mating.

## Methods

### Diet manipulation

We first established our laboratory population of *Peromyscus gossypinus* from wild-caught individuals in 2009 (Delaney and Hoekstra 2018). We maintained a large colony of mice on a standard diet (Purina Iso Pro 5P76) and manipulated diet by providing parents either garlic- or orange-flavored water upon mate pairing: we diluted either 2 μl of Chinese garlic oil or orange oil (both from Sigma Aldrich) into 400 ml of distilled water (0.0005% v/v) and mixed by shaking vigorously – these dilutions did not cause mice to alter their water consumption. We replaced the flavored water every 9-10 days to preserve freshness. Offspring were thus exposed to these chemicals *in utero* (in rodents the olfactory system is functional before birth [Pedersen et al. 1983; Todrank et al. 2011]) and postnatally through weaning, which occurred at 23 days of age. At weaning, we assigned offspring as either a “stimulus” or “chooser”. Stimulus mice were weaned and provided the same flavored water as their parents until their use in trials; chooser mice were weaned and returned to unflavored water.

### Assessment of mate preferences

Using an electronically-controlled gated choice apparatus (Figure 1A; described in Delaney and Hoekstra 2018), we tested the mating preferences of adult mice (> 80 days old) for opposite sex stimulus individuals that were raised on either garlic- or orange-flavored water. We implanted all mice with small radio-frequency identification (RFID) transponders (1.4 mm × 9 mm, ISO FDX-B, Planet ID Gmbh) in the interscapular region. We next programmed antennae to open and close gates in our linear, three-chambered apparatus depending on the identity of a mouse’s RFID: we allowed the designated chooser mouse (i.e. the individual whose preference we tested) to pass freely through all three chambers while constraining two stimulus mice, one each to the left and right cage. We tested individual preferences of 12 to 15 chooser mice from each diet and sex in the gated apparatus for an opposite sex mouse of either the same or alternate diet (Figure 1A). Stimuli mice were fed flavored water up until the start of each trial; during trials, unflavored water was added to all cages with the assumption that the dietary cues, such as odors, from garlic- and orange-fed stimulus mice would persist on the stimulus mice for the duration of the trial.

**Figure 1.**
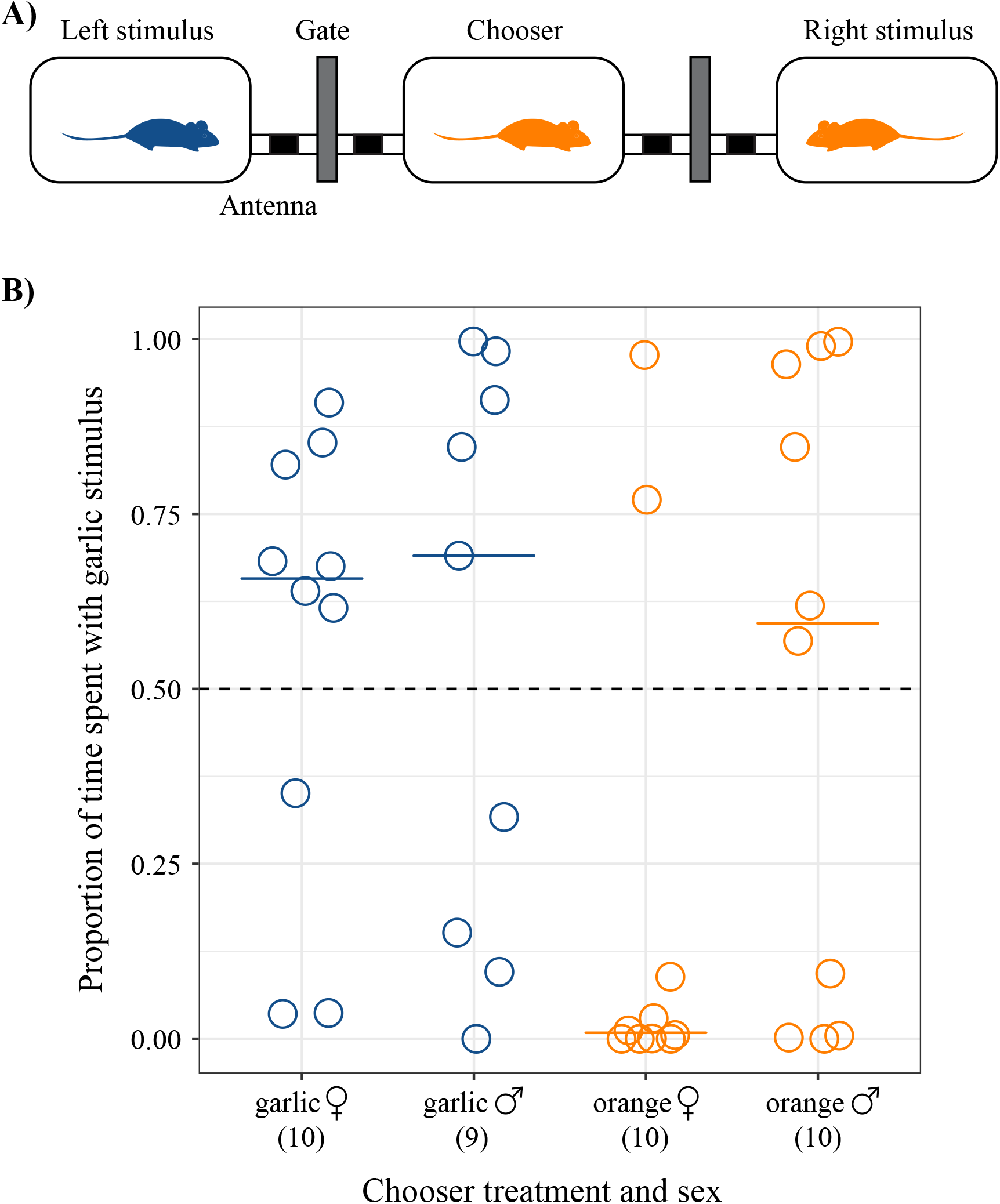
Diet-based assortative mating preferences in female and male *Peromyscus gossypinus*. (A) Schematic of the electronically-controlled gated mate choice apparatus used to measure mating preferences. The apparatus contains three rat cages, each separated by RFID-activated antennae and gates. In the scenario depicted, an orange “chooser” mouse is given a choice between garlic and orange “stimuli” mice of the opposite sex. (B) The dotted line represents equal time with both stimuli: values above the line indicated the garlic stimulus was preferred, and below, the orange stimulus was preferred. Each dot represents the preference of a chooser mouse that was raised with either garlic-fed (blue) or orange-fed (orange) parents. Sample sizes are indicated in parentheses under each treatment group.

For each trial, we added the sexually mature chooser – a virgin female in proestrus/estrus (determined by vaginal lavage) or a virgin male – to the apparatus for one day to acclimate, adding used nesting material from the stimulus mice to the flanking cages. The following day, we added virgin stimulus mice (females were in proestrus or estrus) to the flanking cages to give them 2-4 hours to acclimate before opening the gates at lights out (4:00 pm; 14:10 hour light:dark cycle). We recorded RFID readings at all antennae for approximately 42 hours and calculated mating preference as the proportion of time spent with the garlic-treated stimulus mouse (arbitrarily chosen as the reference) divided by the total time spent with both stimulus mice. We only analyzed trials in which the chooser mouse investigated both cages during the acclimation period, spent at least 10 minutes investigating one or both stimulus mice during the trial, and the stimulus mice were constrained to their cages for > 75% of the trial period.

To assess whether male and female choosers preferred stimuli based on their parental diet, we recorded each chooser’s most preferred stimulus (defined as whichever stimulus the chooser spent more time with). Importantly, we previously showed that the proportion of time a chooser spent with a stimulus in our gated mate-choice apparatus accurately predicts copulation (Delaney and Hoekstra 2018), enabling us to convert chooser preference to a binary variable (garlic mate preferred or orange mate preferred). We used one-sided binomial tests to assess if garlic females, garlic males, orange females, and orange males spent more time with stimuli of the same diet. Additionally, we used a Fisher’s Exact test to determine if preferences for garlic versus orange were significantly different by between females by diet or between males by diet.

### Estimate of sexual isolation attributable to diet

To quantify the amount of reproductive isolation that could arise from diet-based mate choice preferences, we estimated the joint sexual isolation index, *I*_PSI_ (Rolán-Alvarez and Caballero 2000), from our female chooser and male chooser trials separately, as the behavior of the stimuli could have varied among males versus females. The *I*_PSI_ index compares observed and expected mating pairs (assuming random mating among individuals) among the four possible mating pair types (garlic ♀ × garlic ♂, garlic ♀ × orange ♂, orange ♀ × garlic ♂, and orange ♀ × orange ♂). A value of −1 indicates that all mating occurred between diet types, +1 indicates that all mating occurred within diet types, and 0 indicates equal pairing among all four mating pair types. We recorded “mating pairs” based on each chooser’s most preferred stimulus (defined as whichever stimulus the chooser spent more time with) and estimated the sexual isolation index in JMATING v. 1.0.8 using these values (Carvajal-Rodriguez & Rolán-Alvarez 2006). We used 10,000 bootstrap replicates to estimate the isolation indices, their standard deviation, and to test the hypothesis that our sexual isolation estimate deviates significantly from random mating (I_PSI_ = 0).

## Results

One-sided binomial tests (assuming that garlic females and males would spend greater time with garlic stimuli mice, and orange females and males would spend greater time with orange stimuli) failed to reject a null hypothesis of random mating preferences (Figure 1B). Only slightly more than half of males preferred their parental diet (5 out of 9 garlic males preferred garlic females; 6 out of 10 orange males preferred orange females); however females were more biased toward mates of the same diet (7 out of 10 garlic females preferred garlic males; 8 out of 10 orange females preferred orange males). When we analyzed the behavioral results by sex, orange female preferences for orange males were marginally significant in the binomial test (*p* = 0.055), but analyzing female preferences by diet or male preferences by diet were non-significant (Fisher’s Exact test, females: *p* = 0.069, males: *p* = 0.656). Nonetheless, the modest biases of females observed in these assays may still contribute to reproductive isolation.

When estimating the sexual isolation index (*I*_PSI_), we found evidence for diet-based assortative mating in females. In combination, orange and garlic female preferences are predicted to generate significant sexual isolation (*I*_PSI_ = 0.52, sd = 0.20, *p* = 0.023), whereas orange and garlic male preferences do not (*I*_PSI_ = −0.05, sd = 0.26, *p* = 0.859). These data indicate that female preferences, while modest, can generate a strong pattern of diet-based assortative mating.

## Discussion

In this study, we manipulated parental diet in *Peromyscus gossypinus* to test the hypothesis that diet-based assortative mating could form via sexual imprinting. Despite small sample sizes, we found that females had a modest preference for males who fed on the same diet as those females’ parents. Because female chooser mice were exposed to garlic and orange-diet cues only in the nest, we can attribute any biased diet-based assortative preferences to sexual imprinting on parental diet. We also showed that these preferences should produce appreciable positive assortment (*I*_PSI_ = 0.52). This level of sexual isolation between garlic- and orange-fed mice would reduce gene flow to a similar extent as that reported between incipient walking stick species (*I*_PSI_ = 0.24-0.53; Nosil et al. 2013) or Nicaraguan cichlid gold and normal morphs (*I*_PSI_ = 0.39; Elmer et al. 2009), and approaching that reported between distinct species, *P. gossypinus* and *P. leucopus* (*I*_PSI_ = 0.65; Delaney and Hoekstra 2018).

While dietary information was learnable and led to modest assortative mating preferences in females, male preferences appeared random with respect to diet. This sex difference was surprising as we previously established that both *P. gossypinus* sexes can sexually imprint on their parents in a cross-fostering experiment with *P. leucopus* (Delaney and Hoekstra 2018). Thus, males are capable of sexual imprinting but in this study either failed to imprint on diet or imprinted on diet but relied more heavily on other cues (e.g. visual, vocal, or chemical cues) to select mates (Rosenthal 2017).

Although our data showed only weak assortative female preferences for diet, our results agree with studies from other mammalian species (e.g. rats, spiny mice, European rabbits, and humans) that have demonstrated that offspring can learn diet cues from their mothers and later exhibit preferences for those learned foods (Galef and Henderson 1972; Porter and Doane 1977; Hepper 1987; Sullivan et al. 1990; Altbackek and Bilko 1995; Schaal et al. 2000). In our study, diet-induced changes in milk (Désage et al. 1996), amniotic fluid (Mennella et al. 1995), and bodily fluids such saliva, urine or feces (Spiegelhalder et al. 1976) may have served as cues for imprinting. Indeed, mammalian chemosensory systems appear to be active *in utero* (Schaal and Orgeur 1992), raising the possibility that dietary learning could even begin before birth. In support of this view, Todrank et al. (2011) found that mice whose mothers ate cherry- or mint-flavored chow pellets developed larger glomeruli in the olfactory bulb and displayed greater sensitivity to detecting these odorants (Todrank et al. 2011). This enhanced chemosensory sensitivity to maternal diet might contribute to the observed sexual imprinting.

While we found that imprinting is a viable mechanism for diet-based assortative mating in rodents, it is unclear if this same mechanism can explain previous observations in other species. For example, appreciable positive assortative mating by diet in threespine stickleback cannot be explained by spatial co-segregation and microhabitat preferences alone (Snowberg and Bolnick 2012; Ingram et al. 2015). Is there a role for imprinting? Three pieces of evidence suggest the possibility: (1) diet alters gut microbiota in stickleback (Bolnick et al. 2014b, a); (2) such alteration is presumably detectable to the fish, as it was previously demonstrated that changes in diet are sufficient to cause diet-based assortative shoaling behavior (Ward et al. 2004); and (3) stickleback sexually imprint and choose mates using paternal olfactory cues (Kozak et al. 2011). Thus, learned preference for diet-derived olfactory traits might provide a mechanistic basis for diet-based assortative mating in stickleback fishes as well.

Overall, our manipulative experiment suggests an important role for sexual imprinting and learned mating preferences in speciation. In the absence of genetic differences, changes in diet caused by divergent natural selection could lead to sexual isolation. Any change that prompts individuals to diverge in diet – for example, through intra- or interspecific competition over limited food – a learning mechanism such as sexual imprinting would easily couple ecological selection with reproductive isolation, allowing for the coexistence of incipient (or even well-diverged) species in sympatry.

## Data accessibility

Behavioral data are provided in the electronic supplementary material.

## Author’s contributions

EKD and HEH designed the project. EKD collected and analyzed the data. EKD and HEH wrote the paper.

## Acknowledgements

We thank N. Chen, K.G. Ferris, and Y.E. Stuart for comments on the manuscript

